# Potency and selectivity of a novel pan-RAS inhibitor in 3D bioprinted organoid tumor models

**DOI:** 10.1101/2025.02.25.640132

**Authors:** Daniela De Nobrega, Logan C. Eiler, Parmanand Ahirwar, Urvi P. Rawal, Chelsea L. Crawford, Donald J. Buchsbaum, Adam B. Keeton, Yulia Y. Maxuitenko, Xi Chen, Gary A. Piazza, Allan Tsung, Karim I. Budhwani

## Abstract

**Background:** Colorectal cancer (CRC) remains a significant global health burden, with KRAS mutations driving ∼40% of cases. Efficacy of recently approved, mutant-specific KRAS inhibitors is limited by intrinsic and adaptive resistance mechanisms. Pan-RAS inhibitors, such as ADT-007, offer broader therapeutic potential by targeting multiple RAS isoforms. Here, we evaluate ADT-007 in 3D bioprinted organoid tumors (BOTs) generated from KRAS-mutant and RAS wild-type (WT) CRC cell lines.

**Methods:** Potency and selectivity of ADT-007 were compared to bortezomib, a proteasome inhibitor, and YM155, a survivin inhibitor, using high-content imaging and ATP-based luminescence assays. Mechanistic studies assessed impact on RAS activation and downstream signaling.

**Results:** ADT-007 exhibited high potency and selectivity in KRAS-mutant BOTs, reducing tumor burdens >30% at nanomolar concentrations, and demonstrated superior selectivity over bortezomib and YM155 with minimal cytotoxicity in RAS-WT BOTs. Mechanistic analysis confirmed ADT-007 inhibited RAS activation and downstream signaling, leading to selective apoptosis induction in KRAS-mutant CRC cells.

**Conclusions:** The selective potency and specificity of ADT-007 warrants further investigation of pan-RAS inhibitors for treating RAS-driven cancers. This study also underscores the translational utility of 3D BOT models for preclinical drug response assessment. Further validation in patient-derived BOTs is necessary to evaluate potential of ADT-007 in clinical settings.

## 1. INTRODUCTION

Colorectal cancer (CRC) remains one of the most common and lethal cancers in the world with an estimated 1.9 million new cases and nearly 1 million deaths in 2022.^1^ Although CRC is more common in high income countries, recent studies point to increasing incidence in low-and middle-income countries.^2^ Despite recent breakthroughs in cancer research and translational medicine, the prognosis for patients with advanced stage CRC remains poor, particularly for patients harboring KRAS mutations. The RAS family of membrane-bound small guanine nucleotide-binding proteins plays a crucial role in transducing signals from extracellular growth factors intracellularly to regulate proliferation, survival, and differentiation of both normal and cancerous cells.^3,4^ Mutations in RAS result in the constitutive activation of RAS isozymes, KRAS, HRAS, or NRAS, which in turn activate downstream signaling pathways, such as RAF-MEK-ERK and PI3K-AKT-mTOR, that promote malignant transformation, disease progression, and metastasis.^5,6^ KRAS mutations are responsible for approximately 25% of all human cancers and about 40% of CRC cases.^7,8^ Multiple KRAS mutations occur in CRC, including G12D (34%), G12V (21%), G13D (20%), G12C (8%), and others (18%).^9^ Given the critical role of KRAS in oncogenesis, the development of new drugs targeting KRAS has been a focal point of hundreds of cancer research labs around the world given their potential for selective killing of cancer cells with mutant KRAS. However, KRAS was long considered undruggable^10^ due to its high affinity for GTP/GDP and the lack of deep pockets for small molecule binding other than the nucleotide binding domain.^11^ The challenges in directly targeting RAS spurred the development of alternative strategies that selectively induce apoptosis, such as inhibiting downstream signaling pathways, although none have significantly improved patient survival.^12^

Meanwhile researchers continue to investigate other therapeutic approaches for treating CRC, including proteasome inhibitors such as bortezomib, which disrupt protein homeostasis and induce apoptosis.^13^ The proteasome, a multi-catalytic proteinase complex, is responsible for degrading ubiquitinated proteins and regulating various cellular processes, including cell cycle progression and survival. The ubiquitin-proteasome proteolytic pathway is implicated in regulating key proteins involved in cell cycle progression and major transcription factors such as p53, nuclear factor-κB (NFκB), and hypoxia-inducible factor-1 (HIF-1).^14^ Inhibition of the proteasome disrupts these processes, leading to the accumulation of pro-apoptotic factors and induction of cell death. Bortezomib, [(1R)-3-methyl-1-[[(2S)-1-oxo-3-phenyl-2[(pyrazinylcarbonyl) amino]propyl]amino]butyl] boronic acid, specifically and selectively inhibits the 26S proteasome, comprising a 20S core and a 19S regulatory complex. By stabilizing the inhibitory molecule IκB, proteasome inhibitors induce apoptosis to suppress cancer progression and metastasis.^13^ Bortezomib has demonstrated significant antitumor activity in preclinical studies across a wide range of cancers, including CRC.^14,15^ Clinical trials have further investigated bortezomib in patients with metastatic CRC, and demonstrated modest antitumor activity. These studies examined both clinical outcomes and molecular mechanisms through tumor biopsy analysis, revealing differential effects on HIF-1α and its transcriptional target, carbonic anhydrase IX (CAIX), indicating potential disruption of response to tumor hypoxia following bortezomib treatment.^13^

Survivin inhibitors such as YM155 have also emerged as another class of apoptosis inducers with potential therapeutic benefits for CRC. Survivin, a member of the inhibitor of apoptosis protein (IAP) family, is highly expressed in most cancers, including CRC, where it plays dual compounding pathogenic roles by promoting cell proliferation and inhibiting apoptosis.^16^ Survivin has been implicated in both chemoresistance and mortality among CRC patients,^17,18^ particularly in CD133+ cancer cells as demonstrated in studies using the CRC cell line, HT29.^19^ Moreover, the interaction between survivin and CD133 may contribute to disease progression.^19,20^ Targeting survivin has thus garnered considerable interest as a therapeutic strategy with small-molecule inhibitors and antisense oligonucleotides being developed to inhibit survivin function and expression. YM155, 1-(2-Methoxyethyl)-2-methyl-4, 9-dioxo-3-(pyrazin-2-ylmethyl)-4, 9-dihydro-1H-naphtho [2, 3-d] imidazolium bromide, is among the small molecule survivin inhibitors investigated for its potential to treat CRC.^21,22^ However, broad expression of survivin in both cancerous and normal cells raises concerns regarding cancer cell selectivity and associated side effects.^23,24^

Although KRAS was historically considered to be undruggable,^10^ recent advances in technology and drug discovery have made possible the development of KRAS^G12C^ inhibitors, such as sotorasib and adagrasib, which can bind to its inactive GDP-bound state, blocking its activation. Mutant-specific KRAS inhibitors have shown promise not only in preclinical studies but also in clinical trials for non-small cell lung cancer and CRC.^25,26^ However, two serious limitations persist due to the allele-specific nature of these inhibitors: (1) the G12C KRAS mutation accounts for only 3% of CRC patients^27^ thereby severely limiting clinical utility, and (2) experimental studies have identified multiple resistance mechanisms to allele-specific KRAS inhibitors including emergence of new KRAS mutations (e.g. Y96C), KRAS amplification, unchecked activation of co-expressed WT RAS isozymes (e.g. via EGFR stimulation), mutations in NRAS, or acquired bypass mechanisms (MET amplification or new oncogenic fusions).^28–32^ This dual constraint of limited clinical utility and potential for multiple resistance mechanisms underscores the urgent need for pan-RAS inhibitors that target a broader range of RAS mutations that would offer therapeutic benefits to a larger patient population and with reduced potential for resistance.

In recent years, the landscape of KRAS-targeted therapies has evolved significantly with the emergence of alternative targeting strategies and a vast array of novel small-molecule inhibitors. Given the heterogeneity of KRAS mutations that drive CRC, a pan-RAS inhibitor would be expected to have a broader scope of therapeutic use and reach to escape both intrinsic and adaptive mechanisms of resistance. Pan-RAS inhibitors, which target multiple RAS mutations and isoforms simultaneously, have shown promise in preclinical models.^33,34^ Despite the long-held belief that pan-RAS inhibitors would be overtly toxic, one such pan-RAS inhibitor, RMC-6236, is currently in Phase III clinical trials,^34^ while another, ADT-1004, an orally bioavailable prodrug of ADT-007, is in preclinical development.^35^

Concurrently, advances in additive manufacturing and biomedical engineering^36^ have led to the development of innovative tools and technologies that enhance our ability to evaluate these and other new target-directed anticancer agents. For instance, emerging *ex vivo* cancer models now incorporate more physiologically relevant 3D bioprinted organoid tumors (BOTs)^37,38^ instead of traditional 2D monolayer cultures that are less predictive of clinical efficacy.^39–41^ BOTs have the potential to be more predictive of clinical efficacy by more closely mimicking the 3D tumor microenvironment, allowing for the investigation of drug efficacy in a setting that better recapitulates the complexity of human tumors.

Herein, we evaluate ADT-007, a novel pan-RAS inhibitor and compare its potency and selectivity to other apoptosis inducers, bortezomib and YM155, for killing KRAS mutant cancer cells using 3D BOTs derived from KRAS-mutant (HCT-116) and WT RAS (HT29) CRC cells. By elucidating the differential potency and selectivity of ADT-007, we aim to accelerate the development of more efficacious and precision treatment strategies for patients with KRAS-mutant CRC. Further, findings from this study will highlight the potential of emerging 3D bioprinted organoid models as an *ex vivo* assay in preclinical evaluation of targeted-directed experimental anticancer drugs.

## 2. RESULTS

### 2.1. RAS selective inhibition of CRC cell growth by ADT-007

The potency and selectivity of ADT-007 to inhibit CRC cell growth was initially determined using two KRAS mutant and two RAS WT human CRC cell lines grown in monolayer culture following three days of treatment and measured based on viable cell number (ATP content). The relative level of active GTP-bound RAS (GTP-RAS) in each cell line was determined by RBD pulldown assays followed by western blot. Total RAS levels and GAPDH, as a loading control, were measured by western blot. Growth IC_50_ values of ADT-007 for G13D mutant KRAS DLD-1 and HCT-116 lines were 4.7 and 10.1 nM, respectively (**Fig. 1A**). By comparison, growth IC_50_ values of ADT-007 for WT RAS, mutant BRAF, HT29 and COLO 205 lines were 2600 and 2430 nM, respectively. Consistent with the known RAS mutational status of these CRC lines, high levels of GTP-RAS and total RAS were detected in DLD-1 and HCT116 compared with HT29 and COLO 205 (**Fig. 1B**). In contrast to ADT-007, RAS selective growth inhibitory activity was not observed in the same cell lines treated with Gefitinib, an inhibitor of the EGF receptor immediately upstream of RAS (**Fig. 1C**) or with GW-5074, an inhibitor of the RAF1 kinase immediately downstream of RAS (**Fig. 1D**). Additional experiments revealed that ADT-007 (72 hours treatment) induced apoptosis of KRAS mutant HCT-116 cells (**Fig. 2A**), but not RAS WT HT29 cells (**Fig. 2B**) as measured by flow cytometry using Annexin V as a biochemical marker of apoptosis.

**Figure 1.**
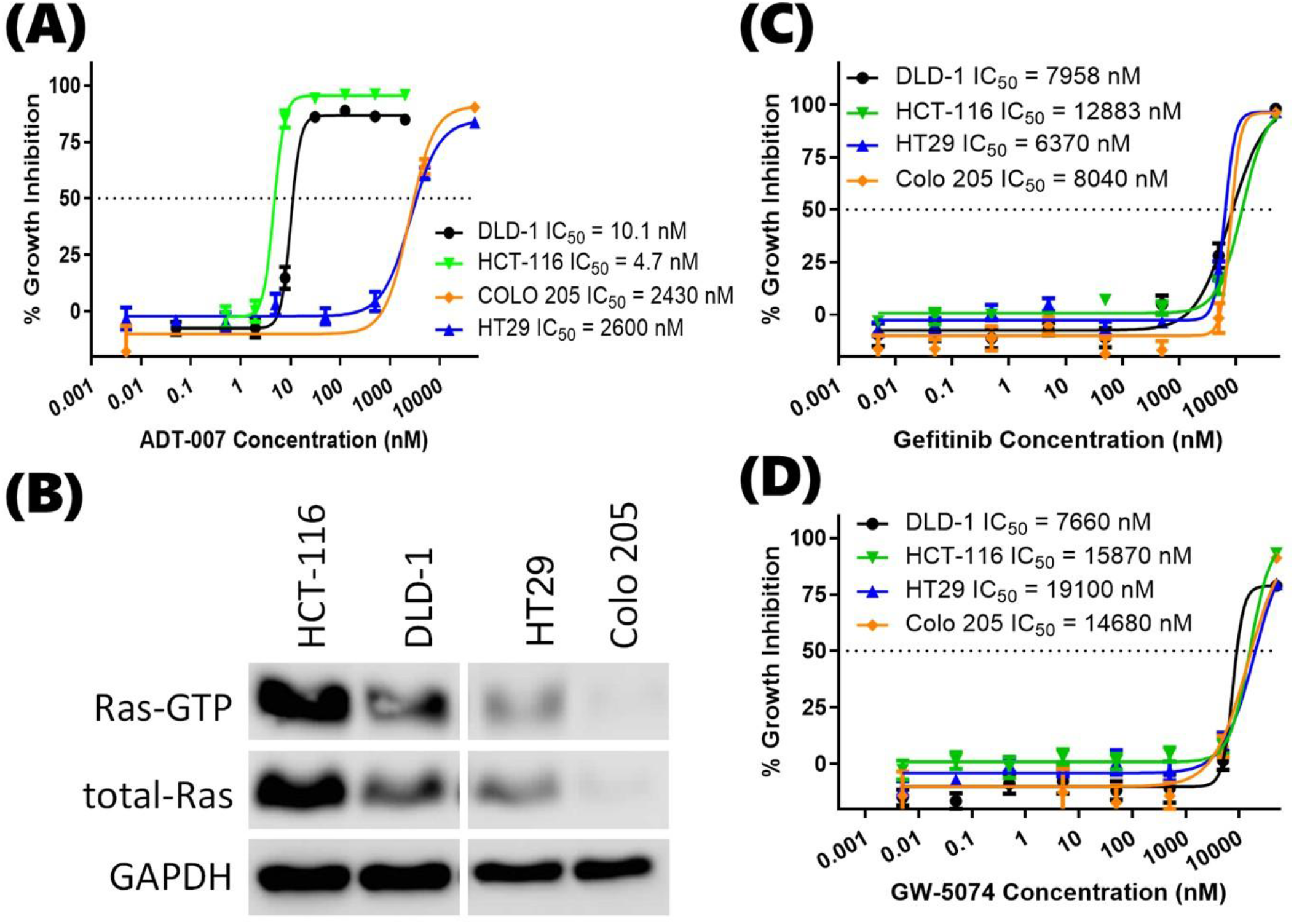
Growth inhibitory potency and RAS selectivity of ADT-007 in standard 2D monolayer cultures of RAS mutant and WT CRC cell lines. **(A)** Dose-dependent growth inhibition of KRAS mutant (DLD-1, HCT-116) versus RAS WT (HT29, COLO-205) cell lines by ADT-007 as determined by cell viability assays following 72 hours of treatment (Promega CellTiter Glo assay). **(B)** Activated and total RAS levels in KRAS mutant (DLD-1, HCT-116) versus RAS WT (HT29, COLO-205) cell lines. **(C)** Dose-dependent growth inhibition of KRAS mutant (DLD-1, HCT-116) versus RAS WT (HT29, COLO-205) cell lines by EGFR inhibitor, Gefitinib. **(D)** Dose-dependent growth inhibition of KRAS mutant (DLD-1, HCT-116) versus RAS WT (HT29, COLO-205) cell lines by RAF1 inhibitor, GW-5074.

**Figure 2.**
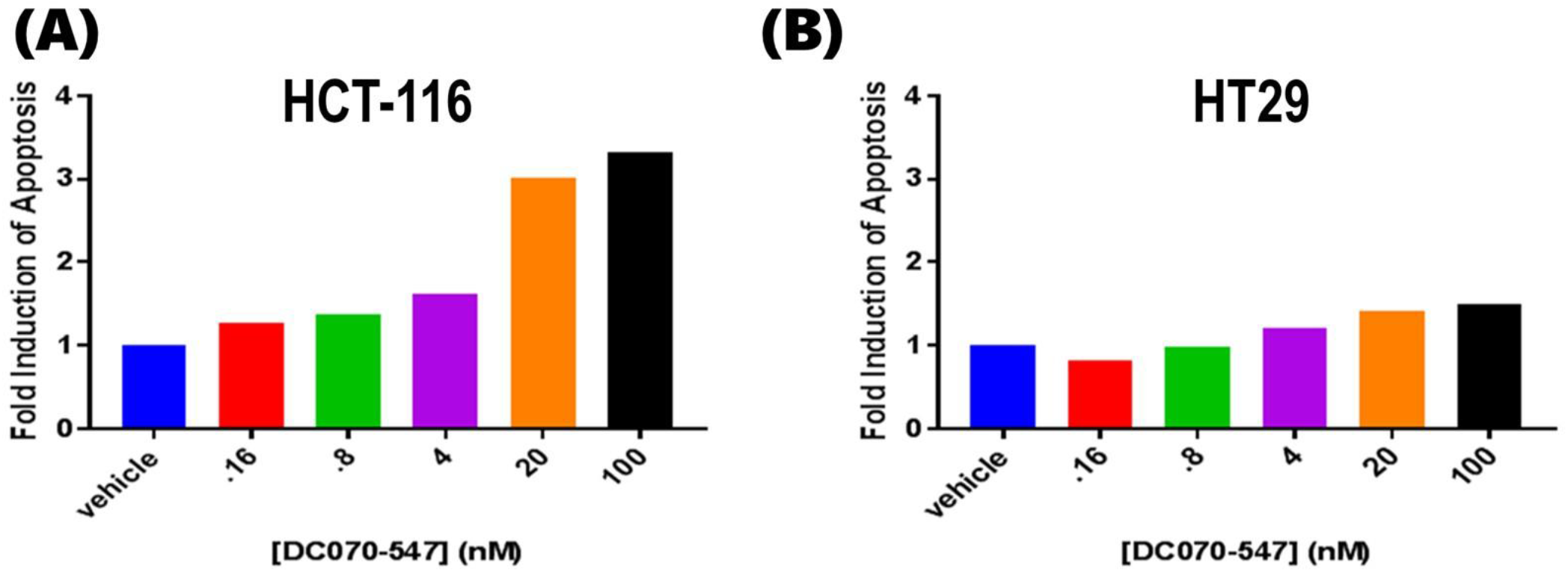
RAS selective apoptosis induction by ADT-007. **(A)** ADT-007 induces apoptosis of KRAS mutant HCT-116 cells as measured by Annexin V levels. **(B)** ADT-007 treatment of RAS WT HT29 cells showed no effect on apoptosis. Cells were plated in 6-well dishes and treated with vehicle (0.1% DMSO) or ADT-007 at the indicated concentration for 72 hours before staining with propidium iodide and Annexin V and analysis by flow cytometry.

We next determined if ADT-007 inhibits RAS activation by measuring GTP-RAS levels with RBD pulldown assays and MAPK signaling using a phospho-specific RAF antibody. G13D KRAS mutant HCT-116 cells were treated with ADT-007 (16 hours) followed by EGF (10 minutes) to activate RAS and MAPK signaling prior to cell lysis and detection by western blot. ADT-007 reduced GTP-RAS levels in a concentration-dependent manner without affecting total RAS levels (**Fig. 3A**). Within the same concentration range, ADT-007 also reduced levels of phosphorylated RAF levels that were increased by EGF stimulation. In addition, ADT-007 decreased levels of PD-L1, which is known to be driven by activated RAS signaling. Additional experiments were performed to confirm a direct effect of ADT-007 on RAS inhibition by treating whole cell lysates from KRAS mutant HCT-116 cells or RAS WT HT29 cells expressing a transfected HRAS mutant (G12V) isoform. As observed in experiments performed by treating intact cells, ADT-007 decreased the levels of activated GTP-RAS in a dose-dependent manner, which rule out indirect effects on upstream signaling events (**Fig. 3B**). Together, these results are consistent with the unique selectivity of ADT-007 to kill KRAS mutant cancer cells in association with suppressing activated RAS as recently reported.^35^

**Figure 3.**
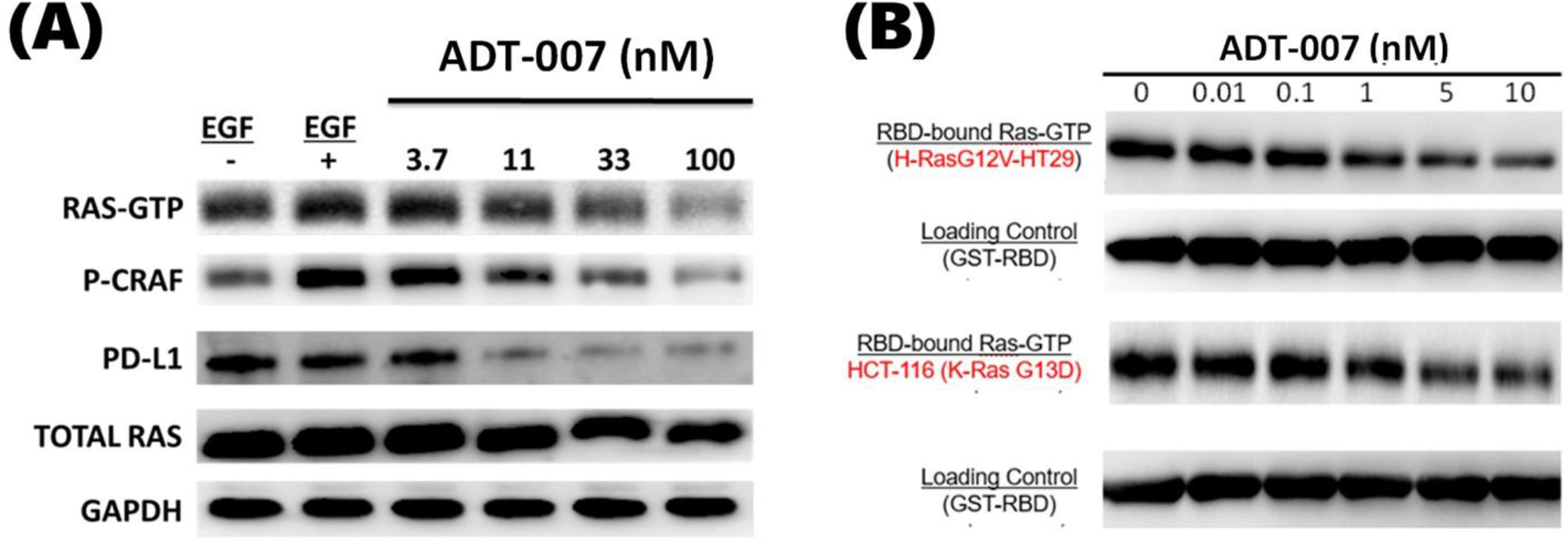
ADT-007 inhibits EGF-induced RAS activation and MAPK signaling. **(A)** Dose-dependent inhibition of EGF-induced GTP-RAS and phospho-CRAF levels by ADT-007 following treatment of KRAS mutant HCT-116 cells. **(B)** Dose-dependent inhibition of GTP-RAS levels by ADT-007 following treatment of lysates from KRAS mutant HCT-116 cells or HT29 cells transfected with mutant G12V HRAS.

### 2.2. Differential response in KRAS mutant CRC BOTS

In prior preclinical studies by other investigators, bortezomib^14,15^ and YM155^21,22^ were reported to inhibit growth of CRC cell lines including KRAS mutant HCT-116 and WT RAS HT29. While both drugs were found to be potent inhibitors, the magnitude of response varied among studies. For instance, studies conducted with HT29 by Pitts et al.^42^ reported an IC_50_ of 500 nM for bortezomib while Suzuki et al.^43^ reported an IC_50_ of 13 nM. Further, since many such studies have been conducted using conventional 2D monolayer cell cultures, we sought to determine growth inhibitory activity for both agents using 3D BOTs that we hypothesize will be more predictive of anticancer activity and compare response with ADT-007, a novel pan-RAS inhibitor. To evaluate anticancer efficacy in our study, a cell population of less than 70% of that in control BOTs at endpoint was used as a proxy for the RECIST criteria of a 30% reduction in tumor burden. This measure reflects the combined effects of growth inhibition and cell death in treated BOTs compared to continued growth observed in control BOTs. In KRAS mutant BOTs, generated with HCT-116 cells, bortezomib was found to be more effective than YM155, which was consistent with differential efficacy profiles reported by other investigators and compiled in the NCI-60 Growth Inhibition Database.^44,45^ Interestingly, for these BOTs, ADT-007 was appreciably more potent than bortezomib and YM155. High-content image analysis of post-treatment BOTs stained with Hoechst 33342 and SYTOX **(Fig. 4A-B)** confirmed that ADT-007 achieved a >30% reduction in tumor burden at lower concentrations (2 nM) compared to either bortezomib (17 nM) or YM155 (17 nM). Metabolic activity, as measured with ATP CellTiter-Glo luminescence assay, was suppressed with lower concentrations of ADT-007 than either bortezomib or YM155, which is consistent with results from high-content imaging. IC_50_ values for ADT-007, bortezomib, and YM155, derived from metabolic activity, were 0.3 nM, 5.8 nM, and 5.4 nM respectively **(Fig. 4C-D)**.

**Figure 4.**
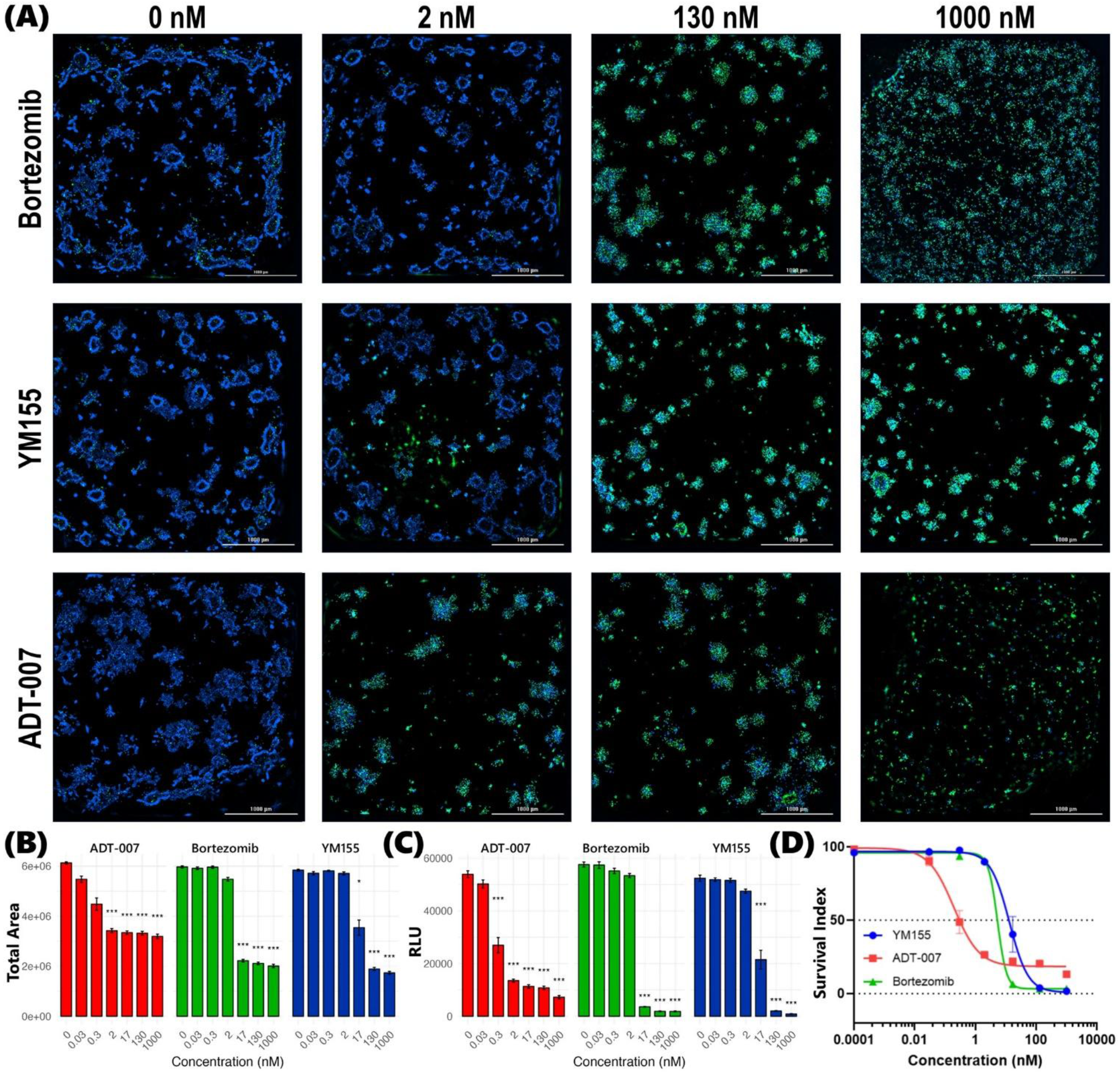
Superior potency of ADT-007 compared to bortezomib and YM155 to inhibit the growth of KRAS-mutant CRC BOTs. **(A)** Representative high-content fluorescence micrographs of nuclear-stained BOTs generated with KRAS-mutant HCT-116 CRC cells. The panels show the effect of treating BOTs with increasing concentrations (left to right) from 0 nM (vehicle) to 1,000 nM of bortezomib (top), YM155 (middle), and ADT-007 (bottom). All images were acquired for multiple z-planes and resolved into single image using a LionHeart imaging system with a 4X objective after treating BOTs for 72h. Blue: Hoechst 33342 (total nuclei); Green: SYTOX Green (dead cells). Scale bar = 1000 µm. **(B)** Dose response relationship between drug concentration and total area occupied by cells derived from high-content imaging analysis. As the drug concentration increases, total cell area decreases for all three drugs. Drug potency was calculated based on reduction in area of Hoechst-positive nuclei and normalized to independent untreated controls. Data points represent mean ± SEM from n = 3 independent experiments, each performed in triplicate. Statistical analysis: *p < 0.05, **p < 0.01, ***p < 0.001 in pairwise t-test against control for each drug concentration. **(C)** Confirmatory dose response from CellTiter-Glo luminescence assay expressed as relative luminescence units (RLU) was obtained in parallel with high-content imaging studies under identical treatment conditions. Data representation and statistical analysis as described above. **(D)** Dose response curves generated from CellTiter-Glo luminescence data using nonlinear regression in GraphPad Prism. Curves were fitted using four-parameter logistic regression. IC_50_ values: ADT-007 = 0.3 nM; bortezomib = 5.8 nM; YM155 = 5.4 nM. *p < 0.01 for ADT-007 vs. bortezomib and YM155.

### 2.3. Differential response in WT RAS CRC BOTs

In WT RAS BOTs generated with HT29 cells, bortezomib was found to be more effective than YM155, which was consistent with differential efficacy profiles reported by other investigators and compiled in the NCI-60 Growth Inhibition Database.^44,45^ While both bortezomib and YM155 showed potency at low concentrations in WT RAS BOTs, ADT-007 was completely inactive, which reflects its unique selectivity for RAS mutant cancer cells. This was evident in both high content image analysis **(Fig. 5A-B)** and ATP CellTiter-Glo luminescence assays **(Fig. 5C-D)**. IC_50_ values for bortezomib and YM155 were 1.6 nM and 7.2 nM, respectively, while an IC_50_ for ADT-007 was greater than 1000 nM.

**Figure 5.**
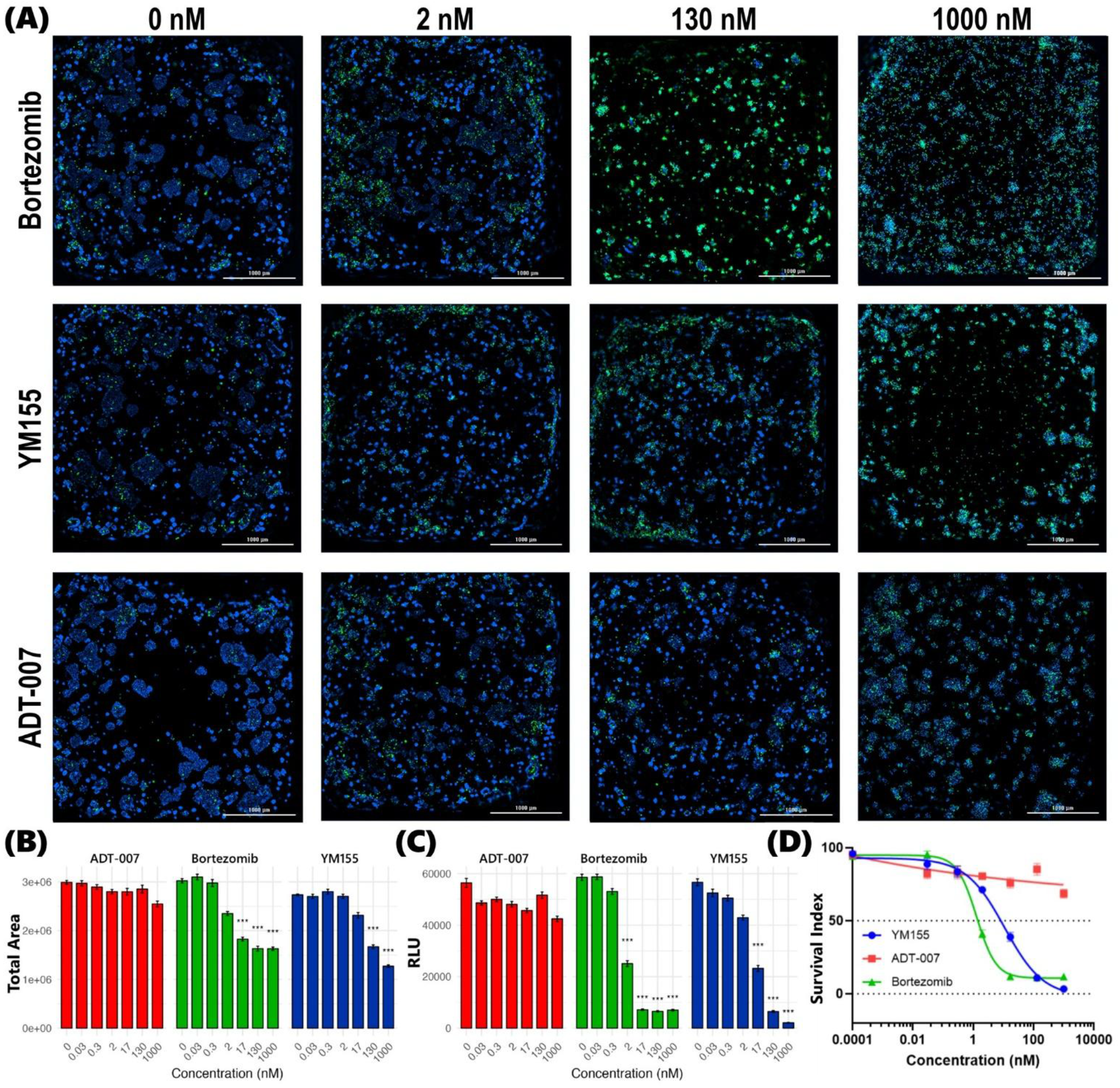
Superior selectivity of ADT-007 compared to bortezomib and YM155 to inhibit the growth of KRAS-mutant CRC BOTs. **(A)** Representative high-content fluorescence micrographs of nuclear-stained BOTs generated with WT RAS HT29 CRC cells. The panels show the effect of treating with increasing concentrations (left to right) from 0 nM (vehicle) to 1,000 nM of bortezomib (top), YM155 (middle), and ADT-007 (bottom). Images acquired using a LionHeart imaging system with a 4X objective after BOTs were treated for 72h. Blue: Hoechst 33342 (total nuclei); Green: SYTOX Green (dead cells). Scale bar = 1000 µm. **(B)** Dose response relationship between drug concentration and total cell area (Hoescht stain). As drug concentration increases for bortezomib and YM155, blue pixel area decreases, indicating an inhibitory effect on cell proliferation. This effect is not observed with ADT-007, as blue area remains nearly constant along the concentration gradient. Drug efficacy was calculated based on reduction in area of Hoechst-positive nuclei and normalized to independent untreated controls. Data points represent mean ± SEM from n = 3 independent experiments, each performed in triplicate. Statistical analysis: *p < 0.05, **p < 0.01, ***p < 0.001 in pairwise t-test against control for each drug concentration. **(C)** Confirmatory dose response from CellTiter-Glo luminescence assay expressed as relative luminescence units (RLU) was obtained in parallel with high-content imaging studies under identical treatment conditions. Data representation and statistical analysis as in (B). **(D)** Dose response curves generated from CellTiter-Glo luminescence data using nonlinear regression in GraphPad Prism. Curves were fitted using four-parameter logistic regression. IC_50_ values: ADT-007 >1000 nM; bortezomib = 1.6 nM; YM155 = 7.2 nM. *p < 0.01 for ADT-007 vs. bortezomib and YM155.

A clear dose dependent response was observed for ADT-007, bortezomib, and YM155 to inhibit the growth of KRAS-mutant CRC BOTs. High-content fluorescence images, captured at multiple z-planes of post-treatment nuclear-stained BOTs, show an increasing proportion of dead cells (SYTOX Green positive) relative to the total number of cells (Hoechst 33342 positive) in tandem with increasing concentrations for each drug. This trend was consistent across all three drugs, albeit with varying degrees of potency. While a similar dose dependent response was observed in WT RAS BOTs for bortezomib and YM155, sensitivity to ADT-007 was at least a hundred-fold lower in these BOTs compared to KRAS-mutant HCT-116 BOTs.

Setting quantitative image analysis aside, qualitative visual comparison of cell death (SYTOX Green positive) captured in **Fig. 4A** (KRAS-mutant HCT-116 BOTs) with that in **Fig. 5A** (WT RAS HT29 BOTs) clearly indicates only a modest effect of ADT-007 in WT RAS BOTs, even at a 130 nM dose, vis-à-vis high potency at only 2 nM in KRAS-mutant BOTs. These observations were quantified and further confirmed by Gen5 image processing algorithms. Specifically, tumor burden decreased with increasing drug concentrations for bortezomib and YM155 in both WT and KRAS-mutant BOTs. However, this trend was only observed for ADT-007 in KRAS-mutant BOTs, resulting in IC_50_ values for ADT-007 of >1000 nM in WT RAS BOTS **(Fig. 5C, 5D)** compared to 0.3 nM in KRAS-mutant BOTs **(Fig. 4C, 4D)**. These results are consistent with previous reports that ADT-007 selectively inhibits mutant RAS by blocking GTP activation of RAS-effector interactions.^33^ Similar dose dependent response profiles were observed in confirmatory CellTiter-Glo luminescence assays. While a significant decrease in luminescence was recorded along the concentration gradient for bortezomib and YM155 in both WT RAS **(Fig. 5C)** and KRAS-mutant BOTs **(Fig. 4C)**, such a trend was limited to KRAS-mutant BOTs for ADT-007 resulting in IC_50_ values that align well with those derived from high-content imaging for all three drugs in both WT RAS and KRAS-mutant BOTs. These results suggest that ADT-007 exhibited superior efficacy in KRAS-mutant HCT-116 BOTs compared to both bortezomib and YM155 as evidenced by lower IC_50_ values in both the high-content imaging cell viability analyses and the confirmatory metabolic activity assay. This enhanced potency of ADT-007 in KRAS-mutant CRC BOTs aligns with its mechanism of action as a pan-RAS inhibitor, directly targeting constitutively activated RAS to block downstream signaling that drives proliferation and survival of these cells. The differential, highly selective response observed for the targeted therapy (ADT-007) compared with the response to both bortezomib and YM155 highlights the need for further development, and demonstrates the promising potential of pan-RAS inhibitors such as ADT-007 in treating RAS-mutant CRC and indeed other RAS-driven cancers.

## 3. DISCUSSION

We recently characterized the ultra-high potency and unique selectivity of ADT-007 to kill cancer cell lines with mutant RAS.^35^ Despite its broad growth inhibitory activity, ADT-007 displayed exquisite target specificity as cancer cells with WT RAS but downstream BRAF mutations, as well as cells from normal tissue, were essentially insensitive to the growth inhibitory activity of ADT-007. The specificity of ADT-007 to kill cancer cells with mutant RAS was attributed to its ability to block GTP activation of RAS and high levels of activated RAS in cancer cells harboring RAS mutations and their dependence on RAS signaling for proliferation and survival, commonly referred to as “RAS addiction”. The requirement of activated RAS for the growth inhibitory activity of ADT-007 was evidenced from experiments shown here where high potency and selectivity to kill CRC cancer cells with mutant KRAS was associated with high levels of activated RAS. We also showed that ADT-007 treatment of such cells decreased GTP-RAS levels and inhibited MAPK signaling. The RAS-selectivity of ADT-007 was not replicated by either upstream or downstream inhibitors of RAS signaling in these cell models. As previously reported and confirmed here, ADT-007 selectively induces apoptosis of KRAS mutant cancer cells, which may be a key advantage over mutant-specific KRAS and other pan-KRAS or pan-RAS inhibitors approved or in development by circumventing resistance.^35^

While initial experiments with ADT-007 were performed in 2D monolayer cultures, there is considerable evidence that 3D tissue cultures are superior to 2D monolayer cell cultures for modeling malignant disease.^39–41^ For example, other investigators have demonstrated greater physiological relevance of 3D cultures to *in vivo* tumorigenicity, metabolic activity, and protein expression compared to that observed in 2D cultures.^46,47^ Thus, in this study we developed and utilized simple 3D BOTs as proxies of WT RAS and KRAS mutant CRC tumors to further study the potency and selectivity of ADT-007 in more predictive preclinical models. Compared to 2D monolayer cultures, the 3D microarchitecture of BOTs can also better replicate potential variations in drug penetration, uptake, cellular response in different regions, and multidimensional influences of drug exposure that would not be readily evident in numerical metabolic activity assays of monolayer cell cultures. For instance, beyond assessing inhibitory or cytotoxic potential of drugs, the approach in this study that combines 3D BOTs with multiplane high-content imaging and a confirmatory conventional metabolic activity ATP assay also enables elucidation of conformational and morphological changes in the microarchitecture of the 3D tumor mimics. Notably, the disruption of characteristic 3D clustering of CRC cells as apparent in multi-plane high-content micrographs of post-treatment BOTs was evident at concentrations appreciably below growth IC_50_ values in 2D monolayer cultures. This high-dimensional data could be utilized as training and testing data for emerging innovative *in silico* drug discovery systems. While this approach may sacrifice speed associated with conventional 2D cell viability assays, the depth and relevance of data obtained could more than compensate for the trade-off. Importantly, our observations from high-content imaging were corroborated by confirmatory metabolic ATP assays, which reported similar dose-dependent trends. This concordance between orthogonal assays strengthens the validity of our findings and underscores the robustness of the 3D BOT model for drug screening. Moreover, the agreement between these distinct methodologies suggests that the observed drug effects are not artifacts of a particular assay.

While our 3D BOT model offers advantages over conventional monolayer cultures, it’s important to note its limitations. BOTs utilized in our study lack key components of the tumor microenvironment, including immune cells, stromal cells, and a complex extracellular matrix. This simplified microenvironment of our BOTs reduces time, complexity, and resources, but it should be noted that this comes at the cost of failing to recapitulate the complex interactions between tumor cells and their surroundings. Additionally, the absence of vasculature in BOTs may affect drug distribution in ways that differ from *in vivo* tumors, potentially impacting our assessment of drug efficacy. It would also be naïve to expect that these findings could be directly applicable *in vivo*, particularly in clinical trials and in the clinic. Factors such as drug metabolism, clearance, and potential off-target effects in other tissues cannot be fully assessed in our simplified 3D BOTs. Despite these limitations, we believe our BOTs provide valuable preclinical insights and demonstrate the potential of high-content multidimensional assays as an effective intermediate step toward more complex patient-derived models. Our study highlights the value of this approach in assessing differential potency and selectivity of ADT-007 in KRAS-mutant versus WT RAS CRC BOTs compared to well-investigated drugs such as bortezomib and YM155. This enhanced performance in KRAS-mutant BOTs, coupled with its reduced cytotoxic effect on WT RAS CRC BOTs, suggests that ADT-007 may offer a more selective approach to targeting cancer cells with mutant RAS, which until recently was considered undruggable. However, it will be crucial to validate these findings in more complex systems, including patient-derived BOTs, which incorporate both tumor and stromal cells from the patient’s own tumor, and eventually in carefully designed clinical trials.

Looking ahead, we recognize the need for further investigation to bridge gaps between these preclinical findings and potential clinical utility. Our planned studies using patient tissue and patient-derived BOTs that incorporate both tumor and stromal cells from the patient’s own tumor, aim to provide more clinically relevant insights into the efficacy and target specificity of ADT-007 and potential advantages over other RAS inhibitors approved or in development. Additionally, investigation into potential resistance mechanisms and combination therapies could further enhance translational potential of ADT-007 and other pan-RAS targeted strategies. In conclusion, while acknowledging the limitations of our current model, this study represents a significant step forward in the preclinical evaluation of targeted therapies for KRAS-mutant CRC. By demonstrating the potential of both ADT-007 and the 3D BOT model, we lay the groundwork for future investigations that could ultimately lead to more effective and personalized treatment strategies for patients with KRAS-mutant CRC and other RAS-driven cancers.

## 4. MATERIALS AND METHODS

### 4.1. Bioprinting organoids

Using previously established bioprinting protocols,^37,38,48^ WT RAS and KRAS mutant 3D BOTs were bioprinted with HT29 and HCT-116 cells, respectively. Both CRC cell lines were acquired from ATCC. HT29 is an established human WT RAS CRC cell line harboring a BRAF^V600E^ mutation with known sensitivity to proteosome and survivin inhibitors. HCT-116 is a KRAS^G13D^ mutant human CRC cell line. Both cell lines were cultured in Dulbecco’s Modified Eagle Media (DMEM) with 10% fetal bovine serum (FBS), 1% Primosin, 1% penicillin-streptomycin at 37°C, 5% CO_2_. Cell viability and counts were assessed using Countess Automated Cell Counter (Vitrogen) with trypan blue. Cell suspensions with <95% cells live were excluded from bioprinting. Bioink components were kept on ice to prevent premature gelation. After printing, BOTs were examined for discoloration, bubbles, or other morphological defects using bright field microscopy, and only defect-free BOTs with uniform cell distribution were utilized for drug screening. 3D BOTs ranged from 300 to 500 µm in thickness. BOTs in clear 384-well plates were used for high-content imaging analysis while those in white 384-well plates were used for metabolic activity measurement with ATP CellTiter-Glo luminescence assay. BOTs were allowed to acclimate for 24 hours before drug treatment. All experiments were conducted in at least three biological replicates and at least three technical replicates. All observations and readouts were also made in triplicates.

### 4.2. Drug treatment

Bortezomib (proteasome inhibitor, MW 384.2) and YM155 (survivin inhibitor, MW 443.3) were acquired from Cayman Chemical at 10 mM in DMSO. ADT-007 (pan-RAS inhibitor, proprietary compound) was provided by Auburn University at 10 mM in DMSO. Each compound was diluted in DMSO and aliquoted at 160 μM to reduce freeze-thaw cycles. Serial dilution and drug administration were performed using a custom adapted epMotion 5070 (Eppendorf) robotic fluid handling system. The maximum concentration of 1000 nM for each drug was freshly prepared before each experiment from a 160 μM stock solution. After dilution, DMSO in all drug solutions was kept below 2% (v/v). Drug solutions were transferred to arrays of sterile PCR tubes for automated dispensing. BOTs were treated with 11 μL of drug at the following concentrations: 0.03 nM, 0.3 nM, 2 nM, 17 nM, 130 nM, 1000 nM. Control BOTs (0 nM) were treated with 11 μL of 0% FBS/DMEM (control: media only, no drug). Following drug treatment, BOTs were incubated under gentle orbital agitation at 37°C, 5% CO_2_ for 72 hours.

### 4.3. High-content imaging

A fluorescent dual-stain cocktail was prepared using Hoechst 33342 (10 μg/mL; Thermo Fisher Scientific, cat. no. H3570) and SYTOX Green (1 μM; Thermo Fisher Scientific, cat. no. S7020) in calcium- and magnesium-free Hank’s Balanced Salt Solution (HBSS; Gibco, cat. no. 14175095). The staining solution was freshly prepared before each experiment and protected from light exposure. 3D BOTs in optically clear 384-well plates (Corning, cat. no. 3712) were stained then incubated at 37°C, 5% CO_2_ under gentle orbital agitation for 2 hours to ensure deep-tissue uniform staining. Automated high-content multi-plane imaging and image processing was performed using a LionHeart Imaging System (BioTek Instruments) with specialized custom protocols designed for analyzing 3D BOTs. Imaging protocol was optimized to capture autofocus z-stack images of Hoechst 33342 (excitation: 377/50 nm, emission: 447/60 nm) and SYTOX Green (excitation: 469/35 nm, emission: 525/39 nm) stained nuclei covering the entire area of each well at each z-plane, ensuring comprehensive data collection for each BOT. Image acquisition parameters, including LED intensity, gain, and integration time, were optimized to maximize signal-to-noise ratio while minimizing phototoxicity and photobleaching. Image analysis was performed using Gen5 imaging software (BioTek Instruments). Total cells in each BOT was approximated by quantifying sum of areas of stained nuclei. Image processing and analysis modules were employed for nuclear segmentation, fluorescence intensity, and sums of object area quantification. Threshold values for Hoechst and SYTOX Green positivity were determined empirically and applied consistently across all analyzed images.

### 4.4. CellTiter GLO

Cell viability was quantified using the CellTiter-Glo Luminescent Cell Viability Assay (Promega, Madison, WI), with modifications to the manufacturer’s protocol. This assay has been shown to be an excellent indicator for cellular activity in medium and high throughput screening of single cell types for its versatility and sensitivity.^49^ The assay exploits the direct correlation between luminescence and cell number over three orders of magnitude, based on the luciferase-catalyzed mono-oxygenation of luciferin in the presence of Mg^2+^, ATP, and molecular oxygen. Thus, the amount of ATP present in the well containing the BOT is directly proportional to the luminescence read from that well, so as the concentration of an active inhibitor increases, luminescence is expected to decrease. After reagent mixture was added to 3D BOTs in white 384-well plates, plates were gently agitated on an orbital shaker at room temperature for 4 minutes to facilitate cell lysis, followed by a 20-minute static incubation in a dark chamber maintained at room temperature to stabilize the luminescent signal. For monolayer cultures, cells were plated at concentration required to achieve 80-90% confluent cultures in black 384-well plates, then treated with the indicated compounds for 72 hours. An equal volume of CellTiter-Glo was added followed by 10 minutes incubation protected from light. Luminescence was quantified using a Biotek Synergy HT plate reader. Cell viability was expressed as a percentage relative to untreated control wells. GraphPad Prism software was used to determine IC_50_ values and generate dose-response curves using logistic regression.

### 4.5 Active RAS pulldown assays

Basal levels of active RAS in a panel of CRC cells were determined using the ThermoFisher Active RAS Pulldown assay kit according to the manufacturer’s protocol. Briefly, sub-confluent cultures of the indicated cell lines in 10 cm dishes were washed with ice cold PBS, and lysates collected in the supplied non-ionic detergent buffer. Protein content was determined by the BioRad DC-assay and normalized to 1 mg/mL. The supplied GST-RBD fusion protein (80 µg) and 15 µL of GSH-agarose were added to 500 µg from each sample, incubated on ice for 1 h. Cell lysates were transferred to spin cups, and GTP-bound RAS captured on agarose was washed thrice by centrifugation. Clarified samples were eluted with Laemmli SDS sample buffer, resolved by PAGE, and transferred to nitrocellulose membrane for western blot analysis with pan-RAS antibody. Companion samples of the lysate were analyzed by western blot analysis for levels of total-RAS and GAPDH as input and loading controls, respectively.

### 4.6. Western blot Analysis

Monolayer cultures were plated in 6-well dishes and allowed to attach before replacement of growth medium with serum-free medium containing vehicle or the indicated concentration of ADT-007 overnight. The following morning, cells were treated for 10 min with either serum free medium alone (EGF-) or 10x concentrated dosing solution of human Epidermal Growth Factor to achieve a final concentration of 30 ng/mL. Growth medium was aspirated, and cells were washed with ice cold PBS. Cells were harvested in RIPA buffer (ThermoFisher) 5 min on ice, then clarified by centrifugation at 4C, 10,000xg. Protein content of supernatants was assayed using the DC protein assay (BioRad) and normalized to achieve 1 mg/mL. Equal samples were resolved by PAGE and electroblotted to nitrocellulose (BioRad). All western blot antibodies were purchased from Cell Signaling Technologies. Blot images were developed using Pierce Supersignal Pico enhanced chemiluminescence reagent in a BioRad Chemidoc imager.

### 4.7. Apoptosis assays

Cell lines were plated in 6-well plates and allowed to grow to 60% confluence. Cells were then incubated with vehicle (0.1% DMSO) or ADT-007 at the indicated concentrations for 72 hours before washing, collection by trypsinization, and staining with propidium iodide/Annexin V according to the kit manufacturer’s protocol (BD Pharmingen, San Diego, CA). Cells were analyzed via flow cytometry using a BD-FACS Canto II (Becton-Dickinson, San Jose, CA, USA). The percentage of apoptotic cells was calculated using DIVA software version 6.1.3 (Becton-Dickinson).

## 5. ACKNOWLEDGMENTS

This work was funded by CerFlux, the National Science Foundation, grant number TI-2321805 and the National Cancer Institute at the National Institutes of Health, grant number 1R43CA254493-01 (Budhwani) and 5R01CA254197-05 (Piazza). We thank our colleagues at The James Cancer Comprehensive Cancer Center at The Ohio State University, The Holden Comprehensive Cancer Center at the University of Iowa, The O’Neal Comprehensive Cancer Center at the University of Alabama at Birmingham, and the Aga Khan University Nairobi Cancer Centre for their support and collaboration. We also thank Drs. Bing Zhu and Antonio Ward for their collaboration and generating experimental data included in this work while at University of South Alabama Mitchell Cancer Institute.

## 6. AUTHOR CONTRIBUTIONS

**DDD**: investigation, data curation, formal analysis, writing - original draft, writing - review & editing. **LCE**: investigation, data curation, formal analysis, writing - original draft, writing - review & editing, visualization. **PA**: investigation, data curation, formal analysis, writing - original draft, writing - review & editing. **UPR**: writing - review & editing. **CLC**: writing - review & editing, project administration. **DJB**: validation. **ABK**: data curation, writing - review & editing, validation. **YYM**: validation. **XC**: resources. **GAP**: writing - review & editing, validation. **AT**: writing - review & editing, validation. **KIB**: conceptualization, methodology, formal analysis, writing - original draft, writing - review & editing, validation, visualization, resources, funding acquisition, supervision. DDD and LCE are co-first authors. All authors have read and agreed to the published version of the manuscript.

## 9. DATA AVAILABILITY

Additional data available upon request.

## 10. CODE AVAILABILITY

Not applicable

